# Acute sleep deprivation induces synaptic remodeling at the soleus muscle neuromuscular junction in rats

**DOI:** 10.1101/2022.06.12.495825

**Authors:** Binney Sharma, Avishek Roy, Trina Sengupta, Lal Chandra Vishwakarma, Anuraag Singh, Ritesh Netam, Tapas Chandra Nag, Nasreen Akhtar, Hruda Nanda Mallick

**Author notes:** Corresponding authors Dr. HN Mallick, Professor, Department of Physiology, Faculty of Medicine & Health Sciences, SGT University, Gurugram, Haryana-122505. B.S. and A.R. contributed equally to this study.

## Abstract

Sleep is important for cognitive and physical performance. Sleep deprivation not only affects neural functions but also results in muscular fatigue. A good night’s sleep reverses these functional derangements caused by sleep deprivation. The role of sleep in brain function has been extensively studied. However, its role in neuromuscular junction or skeletal muscle morphology is sparsely addressed although skeletal muscle atonia and suspended thermoregulation during rapid eye movement sleep possibly provides a conducive environment for the muscle to rest and repair; somewhat similar to slow-wave sleep for synaptic downscaling. In the present study, we have investigated the effect of 24 h sleep deprivation on the neuromuscular junction morphology and neurochemistry using electron microscopy and immunohistochemistry in the rat soleus muscle. Acute sleep deprivation altered synaptic ultra-structure viz. mitochondria, synaptic vesicle, synaptic proteins, basal lamina, and junctional folds needed for neuromuscular transmission. Further acute sleep deprivation showed the depletion of the neurotransmitter acetylcholine and the overactivity of its degrading enzyme acetylcholine esterase at the neuromuscular junction. The impact of sleep deprivation on synaptic homeostasis in the brain has been extensively reported recently. The present evidence from our studies shows new information on the role of sleep on neuromuscular junction homeostasis and its functioning.

**Statement of significance:** Sleep causes synaptic downscaling in the brain, and allows the brain to carry out various housekeeping functions. Here we have reported that the function of the sleep-wake cycle on the synaptic homeostasis extends beyond the brain. Acute sleep deprivation caused significant alteration at ultra and macrostructure of antigravity muscle and the neuromuscular junction along with adaptation to new fiber type in rats. These morpho-functional changes were well correlated with the biochemical assessment of the acetylcholine at the neuromuscular junction. These changes were partially recovered when the rats were allowed to recover from sleep deprivation. The findings suggest a new avenue for a sleep study; employing the neuromuscular junction for exploring the effect of sleep at energy and synaptic homeostasis levels.

## Introduction

Sleep is an enigmatic physiological state, essential for our general and emotional well-being^1^. Most of the studies focus on the association of sleep with the brain. The role of sleep in neurogenesis and synaptic plasticity of the brain is well documented ^2–5^. The slow-wave sleep (SWS) and rapid eye movement (REM) sleep favor neurogenesis in the brain areas like the subventricular zone and dentate gyrus of hippocampal proper, which later proliferates into adult brain neurons^6, 7^. Sleep consolidates memory while sleep deprivation impairs memory consolidation^8^. Studies in *Drosophila* have shown the up and downregulation of many synaptic proteins in the brain during sleep and wake^5^. Deranged cognitive function is associated with sleep deprivation^9^.

During wakefulness, synaptic potentiation is strengthened by glutamatergic synapses. Sleep reduces this energy burden by causing synaptic downscaling, sparing the most robust neural connections^10^. This theory is supported by an experiment where the application of long-term potentiation enhancing neuromodulators caused local cortical slow-wave activity (SWA), and also improved learning of a particular task following sleep^9^. However, as pointed by Matthew Walker in his book “Why We Sleep: The New Science of Sleep and Dreams” - there does not seem to be one major organ in the body or process within the brain that is not optimally enhanced by sleep and detrimentally impaired when we do not get enough sleep^11^. Previous reports suggest that sleep has a restorative effect at the systemic, cellular, and network-level of various organs other than the brain^12–14^. The most consistent daytime complaint associated with insomnia or sleep deprivation is fatigue^15–17^. Sleep is usually acknowledged as an effective countermeasure against fatigue^18, 19^. Decrease in muscle tone as we enter into SWS from the wake, and subsequent muscle atonia during REM sleep is a landmark electrophysiological signature of sleep besides EEG changes^20^. Moreover, REM sleep muscle atonia is because of postsynaptic inhibition of motor neurons^21^.

However, our knowledge of the effect of sleep on muscle as an excitable tissue is limited. The hypothesized effects of sleep loss on muscle function have been largely inferred based on metabolic and endocrine phenomena that accompany long-term REM sleep deprivation^20, 21^. There is paucity of studies on the morphology and physiology of neuromuscular junction (NMJ), after acute sleep deprivation. In this novel study, we have employed the NMJ as a prototype of the peripheral synapse^22, 23^ to investigate the function of sleep on the muscle and the NMJ in rats. We have investigated the histochemical, neuro-morphometric, muscle type adaptation, biochemical estimation of the neurotransmitter, and transmission electron microscopic (TEM) ultra-structure of the synaptic level changes of NMJ during sleep, after 24 h sleep deprivation, and after recovery from 24 h sleep deprivation in the male Wistar rats.

## Methods

### Experimental model and subject details

Adult male Wistar rats (n=30) were used for the experiments. All the experiments were performed following the U.S. National Institute of Health guidelines for the care of animals in research^24^. Efforts were made to minimize the suffering of the animals and also their number. The rats were divided into three groups where the group I rats had a normal sleep-wake (SW) cycle, group II rats were subjected to 24 h sleep deprivation, and group III rats were allowed to recover from 24 h sleep deprivation. The Institutional Animal Ethics Committee approved all the methods used in this research work (960/IAEC/16).

### Surgical implantation of electrodes for sleep recording

The rats were intraperitoneally anesthetized with sodium pentobarbitone (Aldrich Thomas Co, USA) at a dose of 40 mg/kg body weight [BW], and then the surgical implantation of electrodes for sleep recording was done. Two stainless-steel screw electrodes 5 of 2 mm diameter were stereotaxically implanted bilaterally on the skull over the frontal cortex (AP 2.0 mm ML ± 4.0mm at A 9.0) as per De Groot’s Atlas (1959), to record electroencephalogram (EEG)^25^. Two 40 G stainless-steel wires were wound in a loop, and soldered to the uninsulated end of flexible radio wires, to record electromyogram (EMG) and electrooculogram (EOG). Two such loop electrodes were bilaterally fixed on the dorsal nuchal muscles and in the external canthus of the eye respectively for the recordings of EMG and EOG^26^. All these electrodes were then soldered to a socket and the entire head assembly was secured to the skull with three implanted anchoring screws and fixed with dental cement (The Bombay Burmah Trading Corporation Ltd, India).

### Experimental design

After the complete recovery from the surgery, in group I (control, n=10) rats, continuous 24 h recording of SW parameters was done on three alternate days, starting from 10:00 h in the morning till the next morning 10:00 h. The control rats were allowed to sleep in their recording chamber kept in a sound-proof room without any disturbance. In group II, (24 h sleep-deprived), after taking three baseline recordings of normal SW cycle, the rats (n=10) were sleep deprived for a period of 24 h by multiple inverted flowerpot method^27, 28^, during which, SW was continuously recorded. For group III (sleep recovery, n=10); after three baseline SW cycles, the rats were subjected to 24 h sleep deprivation, and then recovery from sleep deprivation was monitored digitally (BIOPAC BSL 4.0 M36 Systems Inc, USA). In all the three groups, a series of experiments were performed including histology of soleus muscle (n=15, 5 each in three groups), myosin ATPase activity of soleus muscle fibers (n=18, 6 each in three groups), immunolabelling for pre and post-synaptic structures at soleus muscle NMJ (n=18, 6 each in three groups), ultrastructural changes of soleus muscle NMJ (n=30, 10 each in three groups), and biochemical estimation of synaptic neurotransmitter. For the neuro-morphometric assessment done by immunolabelling experiment, 240 soleus muscle 6 NMJs was studied via fluorescence imaging, where 80 NMJs were studied for the three groups individually. To evaluate ultrastructural changes in the NMJ via TEM we have used muscle tissue samples from 10 rats for each of the three groups (n=30). Soleus muscle homogenate was used for biochemical estimation of NMJ neurotransmitter acetylcholine (ACh) and its rate-limiting enzyme acetylcholine esterase (AChE).

### Sleep deprivation method

Multiple inverted flowerpot method was used to sleep deprive the group II and III rats. A single rat was placed at one time on an inverted round flowerpot of 6.5 cm diameter kept in a water tank of dimension (123 x 44 x 44) cm. Eight such inverted round flowerpots were kept in a tank filled with water to a level 1 cm below the upper surface of the platforms, where food was kept on each of the platforms. Continuous electrophysiological monitoring was performed in the rat subjected to the sleep deprivation paradigm. After 24 h of sleep deprivation, rats were returned to their home cage where *ad libitum* food and water were provided to them. Further 1 day of sleep recovery recording was continued in their home cage for the group III rats. This multiple platform method is an effective method for sleep deprivation as well as to study the sleep rebound effect^27, 28^.

### Acquisition and quantification of sleep parameters

During one week of postoperative recovery from the surgery, the rats were assessed by observing the body temperature, food-water intake, and general behavior. After recovery from the surgery, the animals were habituated and allowed to move freely in the recording chamber for a day, with the EEG, EOG, and EMG cables connected to the head assembly. The recording chamber was maintained at room temperature and lights-on period same as that maintained in the animal house. Sleep parameters were recorded though a digital polygraph (BIOPAC BSL 4.0 M36 Systems Inc, USA). Continuous 24 h recording of SW stages were collected at an interval of 15 s, and were averaged for a 15 min epoch and are represented as the mean ± standard error of the mean (SEM). Manual scoring of sleep was done for these 15 s epochs and staged as SWS (S) and REM sleep^29^. The SWS was classified as light slow-wave (S1) and deep slow-wave sleep (S2), wakefulness as quiet wake (QW), and active wake (AW).

### Tissue isolation and harvesting

After completion of the sleep deprivation protocol, animals were sacrificed by asphyxiation with CO_2_ and soleus muscle was dissected in dehumidified filter paper placed on ice. This is done to avoid stress response in the tissue of interest. All excisions were done keeping in mind not to cause unnecessary damage to the vasculature and excessive bleeding that would compromise the tissue viability. To assess the muscle mass, the wet weight of the soleus muscle of all the animals from the three groups has been obtained (n=6 each group). For histological preparation, we have placed the dissected muscle directly into isopentane (2-methyl butane, Sigma Aldrich, USA) pre-cooled in liquid nitrogen until processed for sectioning. For histological and whole mount preparations samples were taken on pre-coated slides in gelatine (3% in 1X PBS, w/v), while for TEM, the copper grid was used to take ultrathin sections. For TEM, muscle tissue underwent intermittent immersion fixation with glutaraldehyde (2.5%, in 1X PB, v/v). For immunostaining, the whole-mount preparation was made followed by fixation and teasing in paraformaldehyde (4% in 1X PB at 4°C). For neurobiochemical analysis wet weight of muscle was taken and homogenized in ice-cold phosphorylated buffer saline (1X) and processed further with centrifugation.

### Sectioning of soleus muscle for histology

The soleus muscle of each rat from the 3 groups was isolated and overnight snap-frozen in pre-cooled isopentane in liquid nitrogen. Then 20 µm transverse sections of the frozen soleus muscle were cut by cryotome and stained with hematoxylin and eosin for histological examination.

### Histochemical assay for myosin ATPase

Muscle sections of 10 µm diameter were preincubated in Tris-Calcium buffer (pH=10.2) at 37°C for 20 min. The slides were then washed 3 times with tap water followed by 3 times washing with deionized water. After incubating the slides in Tris-Calcium + ATPase solution (pH=9.4) at 37°C for 30 min, it was twice washed with 1% Calcium Chloride and 2% Cobalt Chloride solution. Slides were then washed with deionized water and were kept in ammonium sulfate solution for 5 to 6 sec. After this, the slides were mounted with gelatin, to quantify myosin ATPase activity.

### Quantification for myosin ATPase activity

The soleus muscle sections (n=6 from each group) were viewed under Nikon upright motorized microscope (Nikon Ni-E with NIS elements, Japan) equipped with fluorescent filters. The cells expressing the myosin fibers were visually identified, and the number of cells was counted in Fiji (Image J) using a cell counter application (http://imagej.nih.ov/ij). Four random fields in all the sections were counted for each rat. All counting values were averaged to give a single value for each rat. The microscopic field under 40x magnifications represented an area of 86056.90 µm^2^.

### Soleus muscle preparation for immunohistochemistry

Soleus muscle samples were obtained from the amputated limb immediately after the animals from each group were sacrificed. Small blocks of tissue, containing full-length muscle fibers from origin to insertion (≈2 cm in length) were removed from each of the selected muscles and were either immediately fixed in 4% paraformaldehyde for 1 h or placed on wet ice. Small bundles of 25-30 muscle fibers were teased out from the larger blocks/whole muscles and of a size suitable for whole-mount preparation. Muscles were immediately fixed in 4% paraformaldehyde for 30 min and then washed in 1% phosphate-buffered saline (PBS). For immunohistochemistry, the selected muscle tissue was cryo-sectioned using the cryostat microtome (MICROM HM 550, Thermo Scientific, USA). It is an open-top, rotary microtome that can section the tissues ranging from 1 to 50 µm in thickness with an increment of 1 µm. Eighty NMJs obtained from each rat of the three groups were then immunolabelled for presynaptic neurofilament with 2H3, for synaptic vesicle proteins with SV2, for synaptophysin with SYP and postsynaptic for ACh receptors (AChR) with α-bungarotoxin (α-BTX).

### Immunolabelling with different markers for pre and post-synaptic proteins

Immunolabelling with α-BTX was done for 30 min followed by application of 4% Triton X for 90 min. The ‘block’ containing 4% bovine serum albumin (BSA) and 2% Triton X was kept for 30 min. The primary antibodies (Mouse anti-2H3, neurofilament 165) at the concentration ratio of 1:50, Mouse anti-SV2 (synaptic vesicles) at the ratio of 1:50 was purchased from Developmental Studies Hybridoma Bank. Mouse anti-Syp (synaptophysin) at the concentration ratio of 1:50 was purchased from Biogene and α-BTX-tetramethyl rhodamine at the concentration ratio of 1:500 was procured from Bungarus multicinctus (Formosan Banded Krait, Sigma Aldrich, USA). All these primary antibodies were added to the block for 3 nights at 4°C and then washed 4 times with 1xPBS. The secondary antibodies, Alexa Fluor 48 donkey anti-mouse IgG secondary antibody at the ratio of 1:50 (Thermo Fischer Scientific) were then added for 1 night at 4°C followed by 4 times washing with 1xPBS. Preparations were then whole-mounted on slides with fluorosheild and DAPI purchased from Sigma Aldrich, USA. These whole mounted slides were then stored at −20°C before imaging.

### Parameters evaluated from NMJ images

For the neuro-morphometric analysis of NMJ obtained after immunohistochemistry in group I, II and III rats, we have used NMJ-morph software^30^. Out of 256 NMJ studied, 80 were analyzed for the three groups of rats. From the 80 NMJ, images obtained from the three groups of rats, core derived, and associated nerve-muscle variables from pre and post-synapse were evaluated. In the pre-synaptic core variables, we have evaluated nerve terminal area and perimeter, the terminal branch number (n), terminal branch point (s), and the total length of the branches (l). At the post-synaptic level, the core variables evaluated were the area, perimeter, and cluster of AChR. The endplate area, perimeter, and diameter were also evaluated at the post-synapse. For the derived variable, at the pre-synaptic level the average branch length and the complexity were evaluated using the following formula:

> Average branch length (µ) = 1/n
>
> Complexity = log10 [no. of terminal branches (n) x no. of branch points (s) x total length of branches (l)] = log10 (n x s x l)

At the post-synaptic level, the average area of the AChR clusters was evaluated. The fragmentation, compactness, overlap, and the area of the synaptic contact were evaluated using the following formula:

> Fragmentation = 1 – [1/ (no. of AChR clusters)]
>
> Compactness = (AChR area/ end plate area) x 100
>
> Overlap = [(total AChR area – unoccupied AChR area) / (total AChR area)] x 100
>
> Area of synaptic contact = total AChR area – the unoccupied area of AChR

For the associated nerve and muscle variable, the diameter of the axon and the muscle fiber along with the number of axonal inputs were determined.

### Soleus muscle preparation for TEM

Soleus muscles from all 3 groups of rats were dissected and immediately fixed with a mixture of 2% (w/v) PFA and 2.5% (w/v) glutaraldehyde (TAAB) in 0.1 M phosphate buffer (PB) pH 7.4 at 4°C for 24 h. Subsequently, samples were post-fixed with 1% (w/v) OsO_4_ supplemented with 1.5% (w/v) potassium ferrocyanide for 1 h on ice, replaced by 1% (w/v) OsO_4_ in sodium cacodylate 0.1 M for an additional hour. Samples were then dehydrated in series of ethanol and infiltrated with propylene oxide (Agar): Durcupan (Agar) (1:1) followed by Durcupan embedding for 48 h. Semi-thin sections were cut (500nm) and stained with toluidine blue to evaluate tissue morphology and select areas for ultrathin sectioning. Ultrathin sections (100 nm) were cut with Ultracut S microtome (Leica), counter-stained with lead citrate, and observed under a Tecnai G2 20 high-resolution transmission electron microscope (Fei Company, The Netherlands) at an operating voltage of 200kV. Images were digitally acquired at 3000-5000 X magnification by a charge-coupled device (CCD) camera using Digital Micrograph software (Gatan, Inc, USA).

### Image acquisition

Photomicrographs were taken with a DS2-Ri2 color camera using NIS Element basic research software (NIKON Instruments, Japan). Fluorescence settings were optimized to achieve the best compromise between image quality and acquisition rate: 8-bit depth, 512 x 512 frame size, x40 magnification, 200±20ms exposure time with 25.6X-32.2X analog gain, and 1 µm z-interval, with manual image acquisition at different depth. Images were acquired with a double filter block of FITC (green; excitation at 475-490nm; bandpass at 483CWL) and TRITC (red; excitation at 545-565nm; band-pass at 555CWL). Fluorescence images were then processed for maximum intensity projections of the z-stacks, using ImageJ software and the Binary Connectivity plugin (downloaded at http://imagej.nih.gov/ij.). For dye-based histological staining, we have used the same camera and software with autofocus function at x40 objective magnification at bright-field filter (color, bandpass 0-∞) and FIJI was used for analysis with 7-8 fields/animals and 4 animals/groups. TEM images were imaged in Tecnai G2-20 with digital micrograph software (Gatan Inc., USA).

### Measurement of neurotransmitter and its rate-limiting enzyme in soleus muscle

To understand the structure-function relationship of the NMJ after 24 h sleep deprivation and recovery we have studied the level of ACh and its rate-limiting enzyme AChE in soleus muscle homogenate. For the same, we have isolated the soleus muscle from 5 animals in each group immediately after the completion of the study by CO_2_ concussion. Further, the sample was snap-frozen in liquid nitrogen and then homogenized with 10% (W/V) ice-cold 0.1M PBS (1X; 7.4 pH) using a homogenizer and emulsifier (Remi Inst.; India) in a customized sample holder. Then this processed sample was centrifuged at 10,000g at 4°C (Hermle, Germany) and the supernatant was aliquoted and preserved in −20°C until further use.

### Acetylcholine in soleus muscle

We have used a colorimetric kit for ACh (Cat no. EACL-100; EnzyChomTM, BioAssay Systems, USA), following the manufacturer protocol where at the final step spectroscopic reading was taken at 570 nm, followed by an incubation within working reagent for 30 min at room temperature. And concentration was calculated from the following formulae:

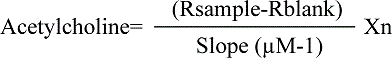

Where R sample and R blank are optical/fluorescence intensity readings of the sample and H_2_O respectively, n is the sample dilution factor.

### Acetylcholinesterase in soleus muscle

We have measured AChE with sandwiched ELISA (Cat no. ER0461; Fine Test, USA) technique. The manufacturer’s protocol was followed for the ELISA and in the final step after the addition of stop solution, optical density was measured at 450 nm and the concentrations of the unknown sample were calculated from the equation obtained from the standard curve drawn out of the six serial dilutions of the standard.

### Statistical analysis

All the data for the various parameters were run though the Shapiro-Wilk normality test. Comparisons of the parametric variables within the three groups were done using one-way analysis of variance (ANOVA) followed by Bonferroni correction. The significant difference between the non-parametric data within the 3 groups was evaluated using the Kruskal-Wallis test followed by Dunn’s post-hoc. Where we did not get any significance, we re-evaluated taking two independent groups individually, and have applied the Mann-Whitney U test. The statistical model used in the present study is following previous literature ^31, 32^. The intact and altered pre-synaptic mitochondrial status in the three groups of rats was analyzed using 2-way ANOVA followed by Bonferroni’s multiple comparison tests. To decipher the role of neurochemical transmission with its structural correlate in NMJ in the 24 h sleep deprived rats, we have performed Spearman’s correlation between ACh and AChE with pre and postsynaptic morphovariables calculated from the electron microscopic examination. All statistical analyses were performed using Graph-Pad Prism 8 software. The significance threshold was set at p<0.05.

## Results

### Sleep-wake cycle during sleep deprivation and recovery sleep

The time spent (%) in the different SW stages of the group I, II, and III rats is depicted in **Fig. 1.A**. There was a significant change in the AW (χ^2^=23.5, p<0.0001), QW (χ^2^=19.7, p<0.0001), S1 (χ^2^=10.2, p=0.002), S2 (χ^2^=23.1, p<0.0001) and REM (χ^2^=19.6, p<0.0001) when compared between the three groups. The control rats spent 54.7 ± 6.3% of their SW time in AW and 1.6±0.5% in QW; while a significant increase in AW was observed in group II rats (p=0.02), which was significantly reduced when given a chance to recover from sleep deprivation (p< 0.0001). A significant increase in QW was seen in the group III when compared to group I (p<0.001) group II (p=0.001). During sleep deprivation, the time spent in S1 and S2 was significantly reduced to 10.0±1.4% (p=0.0003) and 0.8±0.3% (p=0.013) respectively compared to group I. In the group III rats, significantly higher time was spent in S2, compared to group II (p<0.0001) rats. Though the time spent in REM sleep in group II (0.3±0.1%) was less compared to group I rats (2.5±0.8%), it did not reach significant threshold. Time spent in REM sleep (6.5±0.3%) was significantly increased in group III from group I (p=0.04) and II (p<0.0001).

**Figure 1:**
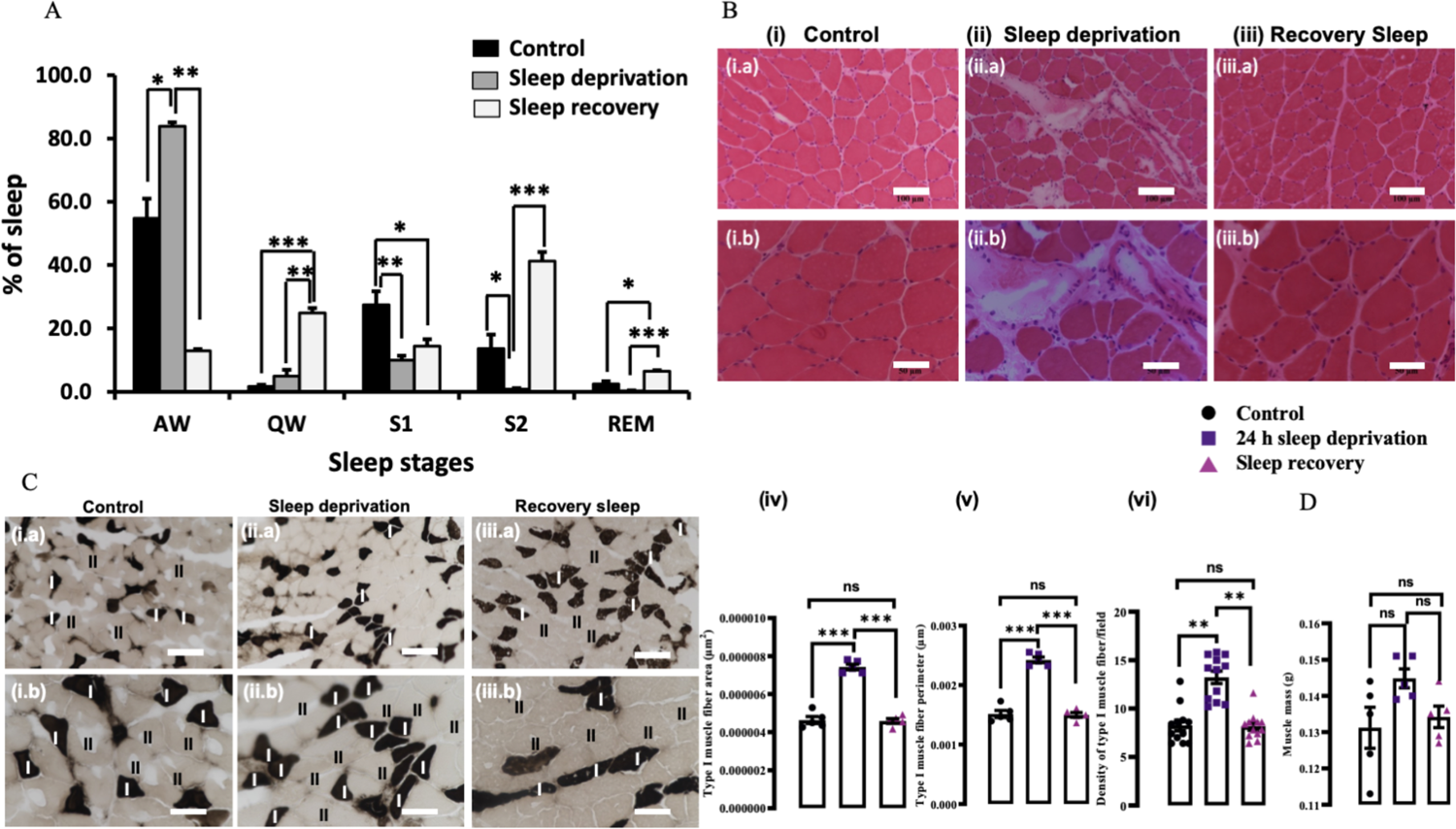
Sleep and sleep deprivation affecting histological changes in soleus muscle of male Wistar rats. (A) Percentage of various sleep stages (%) across the 24 h duration recorded from male Wistar rats (n=30). Recording of sleep-wake parameters during the control, 24 h sleep deprivation, and sleep recovery are shown. The asterisk denotes statistically significant differences (*p<0.05, **p<0.005, ***p<0.0005) between control (▪) sleep deprivation (▪), and sleep recovery (▪) as revealed from the Kruskal-Wallis test followed by Dunn’s multiple comparison test. AW: active wake; QW: quiet wake; S1: light slow-wave sleep; S2: deep slow-wave sleep; REM: rapid eye movement sleep; (B) Haematoxylin and eosin staining for histology of the transverse section of the soleus muscle delineating the changes observed only in sleep deprivation group compared to other in (a) low (20X) and (b) high magnification (40X) in (i) control, (ii) 24 h sleep deprivation, (iii) recovery sleep; (C) Comparative myosin ATPase staining of transverse section of soleus muscle appearing type I fiber darker than type II is marked as ‘I’ and ‘II’ in (i) control, (ii) 24 h sleep deprivation, (iii) recovery sleep with low (20X) and high (40X) magnification; this was further quantified in FIJI showing area, perimeter, density of type I (iv-vi.) fiber in rats with normal sleep (n=6, indicated by ●), 24 h sleep deprivation (n=6, indicated by ▪) and recovery sleep (n=6, indicated by ▴); soleus muscle mass (D.); the statistically significant values are represented as * after applying one-way ANOVA followed by Bonferroni post-hoc test. *** represents p = 0.0006 and **** represents p < 0.0001; data is represented as mean ± SEM showing individual data-points (n=6/ group); scale bar=100µm in 20X and 50µm in 40X.

### Histological changes in soleus muscle

In order to understand the macrostructure of the soleus muscle after 24 h sleep deprivation, we performed hematoxylin and eosin staining on transverse section. In the group I animals, subjective observation of high magnification images show maintenance of fascicular architecture and minimal variation in size of fiber **(Fig. 1.B.i.a-b)**, while in the group II rats there was presence of fibers with myophagocytosis **(Fig. 1.B.ii.a-b)**. Furthermore, mild perimyseal inflammatory infiltrates were also noted in the group II rats. In the group III, we have observed few atrophic and angulated fibers with maintenance of fascicular architecture in the soleus muscle **(Fig. 1.B.iii.a-b)**. These findings suggest a change in morphology of anti-gravity muscle i.e. soleus muscle after 24 h sleep deprivation.

### Histochemical assay for myofibrillar ATPase

The photomicrographic representation of histochemical assay of the soleus muscle type I fiber obtained from the group I, II, and III rats are depicted in **Fig 1.C**. The cell counts were done on the stained images using Fiji (Image J) software (http://imagej.nih.ov/ij)^33^. Histometric analyses revealed an increase in the size and density of type I fibers in the group II compared to group I and III (**Fig 1.C.iv and v**). After 24 h sleep deprivation, there was a significant increase in area and perimeter of type I fibers compared to group I and III (F_(1.1,_ 15 _4.5)_=208.7, p<0.0001). When compared between group II and III, there was a significant decrease in the area and perimeter of cells per filed (p=0.03) as shown in **Fig 1.C. iv and v**. Additionally, we also have found significant increase in density of type I fiber/field examined in group II compared to group I (p=0.0003) and III (p= 0.0005; **Fig 1.C.vi**). No significant changes were observed in wet weight of soleus muscle (**Fig. 1.D)** after 24 h sleep deprivation when compared to the group I and III (χ^2^= 4.63, p= 0.094).

### Immunohistochemistry of neuromuscular junction

The presynaptic protein markers viz. neurofilament, synaptic vesicle, and synaptophysin are immunolabelled with 2H3, 2SV2, SYP respectively (green fluorescence). The postsynaptic AChR is immunolabelled with α-BTX (red fluorescence). In the group I rats, the NMJs appeared as pretzel-like structures (**Fig 2. A.i**). In the group II rats, there were characteristic changes in the neurofilament (**Fig 2.A.ii)**, which is further supported from neuromorphometric analysis (**Fig 3**).

**Figure 2:**
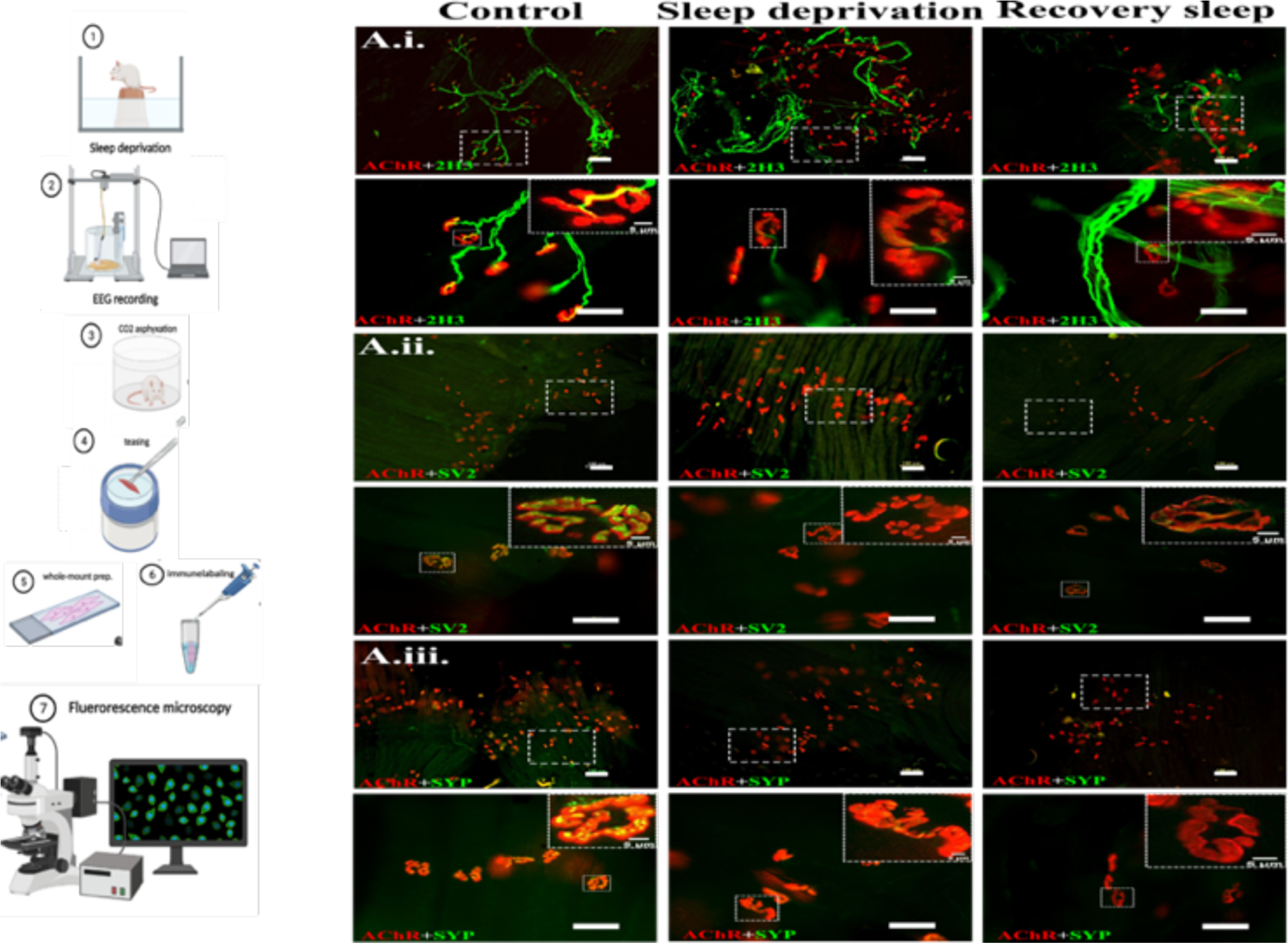
Immunofluorescence images of neuromuscular junctions of three different presynaptic markers colocalized with acetylcholine receptor Whole-mount preparation and imaging work-flow illustration (made using BioRender); (A.i) panel showing ACh+2H3 signals at low and high magnification of control, 24 h sleep deprivation and recovery sleep with zoomed NMJ (a-f.), (A.ii) panel shows ACh+SV2 signals at low and high magnification of control, 24 h sleep deprivation and recovery sleep with zoomed NMJ (a-f.), (A.iii) panel showing ACh+SYP signals at low and high magnification of control, 24 h sleep deprivation and recovery sleep with zoomed NMJ (a-f.) in soleus muscle whole-mount preparations. The postsynaptic marker (ACh) is in red and the presynaptic markers (2H3, SV2, SYP) are in green. Single NMJ is magnified in the inset (scale bar= 5µm) of high magnification images; Scale bar=100µm in low (20X) and 50µm in high (40X) magnification images.

**Figure 3:**
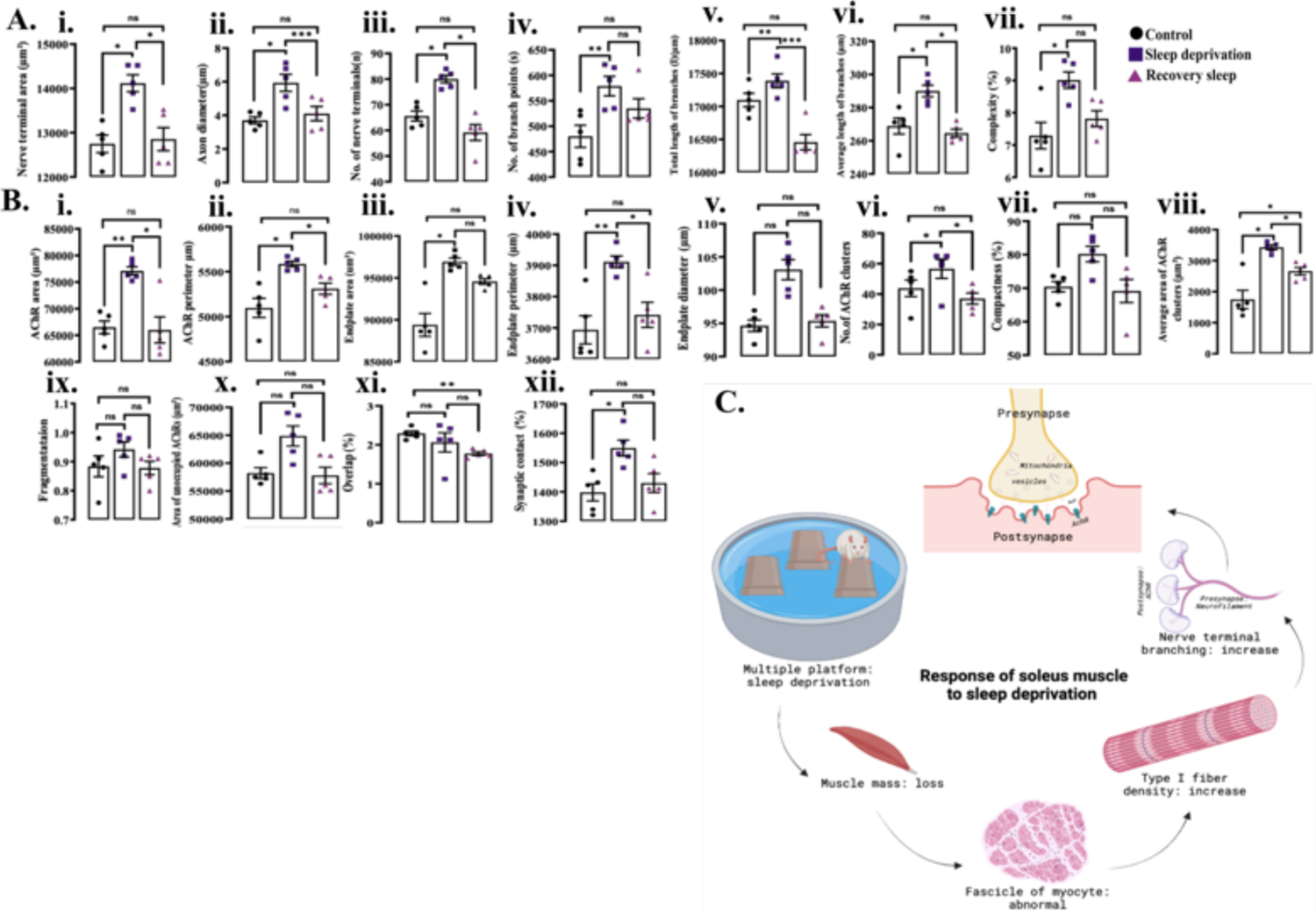
Morphometric measurements in immune co-localization of pre and postsynaptic proteins at the neuromuscular junction of rats after sleep deprivation and recovery sleep. (A) Presynaptic morphometric variables analyzed using ‘NMJ-Morph’ work flow, (i) nerve terminal area (µm^2^), (ii) axon diameter (µm), (iii) number of nerve terminal (n), (iv) number of branch point(s), (v) total length of branches (l in µm), (vi) average branch length (µm), and (vii) complexity (%) in the NMJ of soleus muscle of rats with normal sleep wake cycle (n=6, indicated by ●), rats subjected to 24 h sleep deprivation (n=6, indicated by ▪) and in rats allowed to recover from 24 h sleep deprivation (n=6, indicated by▴); (B) Post synaptic morphometric variables (i) ACh area (µm^2^), (ii) ACh perimeter (µm), (iii) endplate area (µm^2^), (iv) endplate perimeter (µm), (v) endplate diameter (µm), (vi) number of ACh clusters, (vii) synaptic compactness (%), (viii) average area of AChs clusters (µm^2^), (ix) synaptic fragmentation, (x) area of unoccupied ACh (µm^2^), (xi) synaptic overlap (%), and (xii) synaptic contact (%) in the NMJ of soleus muscle of rats with normal sleep wake (n=6, indicated by ●), rats subjected to 24 h sleep deprivation (n=6, indicated by ▪) and in rats allowed to recover from 24 h sleep deprivation (n=6, indicated by▴); (C) Illustration explaining the changes after sleep deprivation in micro and macro structure (made using BioRender); The statistically significant values are represented as * after applying one-way ANOVA followed by Bonferroni post-hoc test for parametric data and Kruskal-Walis with Dunn’s multiple choice test was performed for non-parametric data. *denotes significance where * represents p<0.05 and ** represents p<0.001, *** p<0.0001 and ns: not significant; data is represented as mean ± SEM with individual data points (n=6/ group); ACh= acetylcholine receptor, 2H3= neurofilament-m, SV2= synaptic glycoprotein 2A, SYP= synaptophysin.

At the postsynaptic level of the group II rats, there was an increase in the size of the endplate area of the AChR, and fragmentation of the pretzel-shaped structure of the NMJ (**Fig 2.A**). All these NMJ changes were partially restored after recovery sleep in the group III rats (**Fig 2.C**). At the pre and postsynaptic terminals of the NMJ of group II rats, there were structural and morphological changes in the size of synaptic vesicle when compared to the group I rats (**Fig 2.A**), which is further revealed from the neuro-morphometric analysis **(Fig 3)**.

### Neuro-morphometric changes at the presynaptic terminal of the neuromuscular junction

A significant increase in the nerve terminal area (F_(1.4,5.4)_=19.7, p=0.004), was observed at the presynaptic level of the group II rats when compared to group I (p=0.02). The nerve terminal area was significantly reduced in group III when compared to group II rats (p=0.04) (**Fig 3.A.i**). There was a significant change in the axon diameter when compared between the three groups (F_(1.0,4.3)_=16.4, p=0.01). An increased (p=0.04) axon diameter was observed in the group II rats when compared to the group I (**Fig 3.A.ii**). A significant difference in the number of nerve terminals was observed between the three groups (F_(1.3,5.1)_=18.63, p=0.006). Further, Dunn’s multiple comparison exhibited a significant increase in group II compared to group I (p=0.006) and III (p=0.02) as indicated in **Fig 3.A.iii**. There was a significant increase (p=0.002) in the number of branch points between control and group II rats (**Fig 3.A.iv**). From **Fig 3.A.v** and **vi**, it is evident that the total branch length (χ^2^=10.3, p=0.0005) was increased in group II (p=0.02) compared to group I and III (p=0.008), and the average branch length (χ^2^=9.8, p=0.001) obtained from the group II rats was significantly more compared to the group I (p=0.01) and III (p=0.01) rats. A significant increase (F_(1.6,6.3)_=7.4, p=0.03) in the complexity (%) of NMJ in the group II rats was observed when compared to the control rats (**Fig 3.A.vii**).

### Neuro-morphometric changes at the post-synapse of the NMJ

A significant change in the postsynaptic AChR area (χ^2^=10.8, p=0.0002) was observed between the group I versus II (p=0.01) and group II versus III (p=0.01) (**Fig 3.B.i**). Significant increase in the AChR perimeter (χ^2^=10.7, p=0.0002), endplate area (χ^2^=11.6, p<0.0001), endplate perimeter (χ^2^=9.1, p=0.004) was observed in group II rats when compared to control (**Fig 3.B.ii-v**). Of these parameters, end-plate area and diameter did not show any significant change after sleep recovery when compared to the group I (**Fig 3.B iii, v**). A significant change was found in the average area of AChR cluster (χ^2^=10.8, p=0.0002) and synaptic contact (χ^2^=8.1, p=0.01) when compared between the three groups (**Fig 3.B. viii, xii**). There was a significant decrease (χ^2^=6.3, p=0.003) in the synaptic overlap when compared between the group I and III rats (**Fig 3.B.xi**).

### Ultra-structural changes in the soleus muscle neuromuscular junction

Transmission electron microscopic images of neuromuscular junction showing presynaptic (nerve terminal) and postsynaptic (covered by sarcolemma) in three experimental conditions is shown in **Fig. 4.Ai.-iii**. In the representative images we have shown a normal cristae in the mitochondria present at presynaptic compartment of control group as compared to that of 24 h sleep deprived and recovery sleep where we have observed swollen mitochondria in the presynaptic nerve terminal. In the recovery sleep we have observed glycogen droplets at the pre-synapse. Further, in order to detail these changes in terms of mitochondrial pool, synaptic vesicle pool and junctional folds we have performed morphometry on the images using image J as described previously^34^.

**Figure 4:**
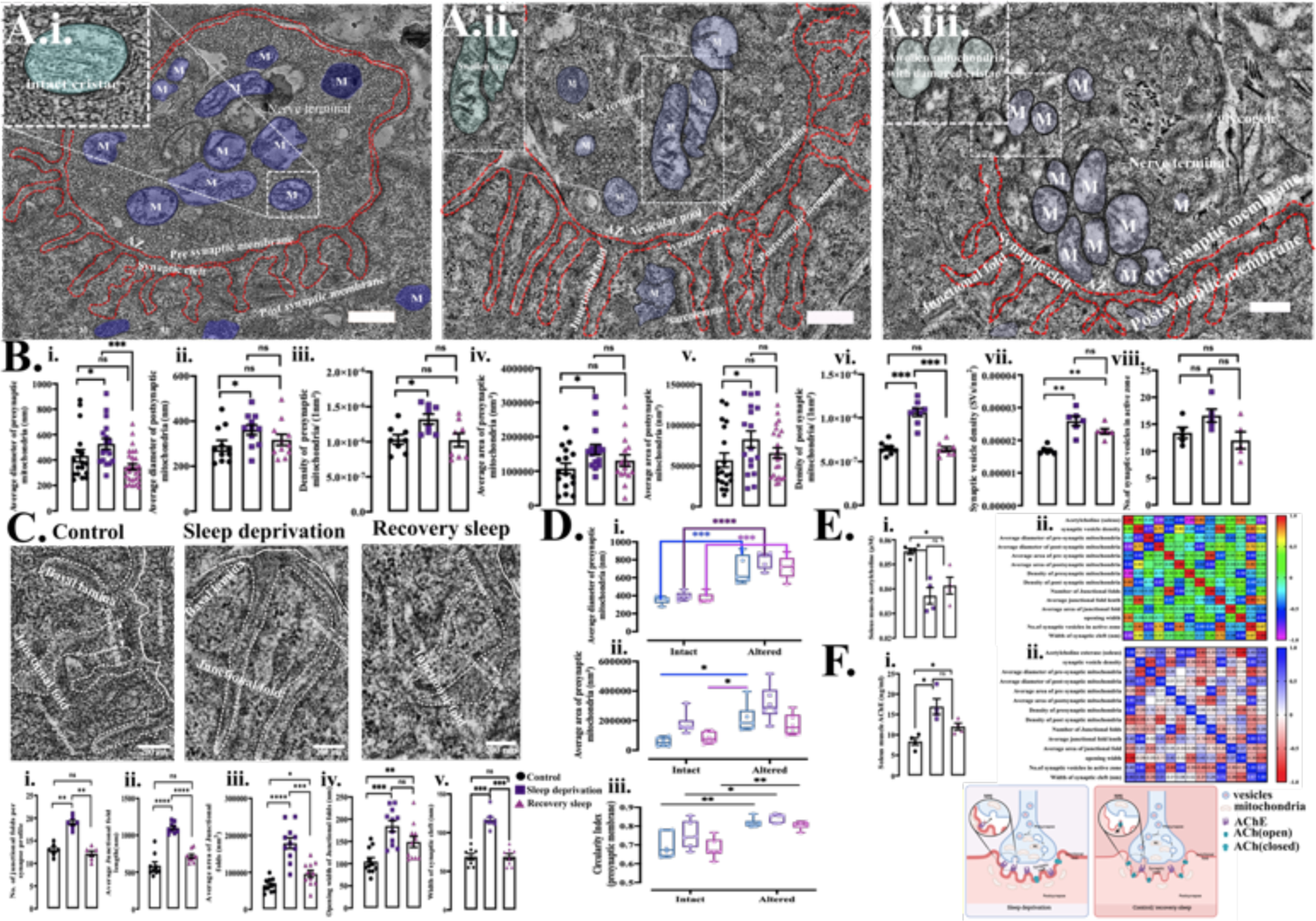
Ultrastructural changes in the transmission electron microscopic examination at the levels of NMJ in the rats after sleep deprivation and recovery sleep Comparative transmission electron microscopic images of (A.i.) control, (A.ii.) 24 h sleep deprivation, (A.iii.) recovery sleep with ultra-structures marked with scribble drawing and mitochondrial changes are magnified in the inset (scale bar= 500nm); (B) Panel showing the morphometric analysis performed in pre and postsynaptic variables average diameter of (i) presynaptic and (ii) postsynaptic terminal mitochondria (nm), the average area of (iii) presynaptic and (iv) postsynaptic terminal mitochondria (nm^2^), the density of (v) presynaptic and (vi) postsynaptic terminal mitochondria per (nm^2^), (vii) total density of synaptic vesicles (SV/nm^2^) in the presynaptic terminal, and the (viii) no. of synaptic vesicles within the 200nm distance of active zone (%); (C) comparative transmission electron microscopic images of the junctional fold at three groups as entitled at the each column with the morphometric analyses (scale bar= 200nm), (i) no. of junctional folds per synapse profile, (ii) junctional fold length (nm), (iii) average area of junctional folds (nm^2^), (iv) opening width of junctional folds (nm), and the (v) width of the synaptic cleft (nm) (%) in the NMJ of soleus muscle of rats with normal sleep wake (n=6, indicated by ●), rats subjected to 24 h sleep deprivation (n=6, indicated by ▪) and in rats allowed to recover from 24 h sleep deprivation (n=6, indicated by▴); (D) Within group comparison between intact versus altered mitochondria, (i) the average diameter of presynaptic terminal mitochondria (nm), (ii) average area of presynaptic terminal mitochondria (nm^2^), (iii) circularity index of mitochondria of the presynaptic terminal in control (blue box-plot), 24 h sleep deprivation (purple box-plot) and sleep recovery (pink box-plot) rats; (E) expression of neurotransmitter and its relationship with the ultra-structural variables, (i) level of acetylcholine in homogenates of soleus muscle, (ii.) correlation matrix between acetylcholine and 13 synaptic variables; (F) expression of rate limiting enzyme in soleus muscle and its relationship with ultra-structural variables, (i.) level of acetylcholinesterase in the soleus muscle homogenate, (ii.) correlation matrix between acetylcholinesterase and 13 synaptic variables. (E) A model reflecting morpho-functional changes due to the 24 h sleep deprivation at the neuromuscular synapse (made using BioRender). The statistically significant values are represented as * after applying one-way ANOVA followed by Bonferroni post-hoc test for parametric data and Kruskal-Walis with Dunn’s multiple-choice test was performed for non-parametric data. *denotes significance where * represents p<0.05, ** represents p<0.001, *** p<0.0001 and ns: not significant; data is represented as mean ± SEM with individual data points (n=6/ group).

### Mitochondrial pool

There was a significant increase (χ^2^=13.02, p<0.0476) in the average diameter of pre-synaptic terminal mitochondria in group II compared to III (p=0.002) (**Fig 4.B.i**). On applying the Mann-Whitney U test, we observed a significant increase in average diameter (U=28, p=0.03) and area (U=98, p=0.04) of post-synaptic mitochondria (**Fig 4.B.ii and v**), and also in the area of pre-synaptic mitochondria (U=70.5, p=0.03) of the group II versus the control rats (**Fig 4.B.iv**). The pre-synaptic (U=9.5, p=0.01) mitochondrial density was increased in group II rats compared to control (**Fig 4.B.iii**). The post-synaptic (χ^2^= 15.6, p=0.0004) mitochondrial density was significantly altered between group I and II (p=0.003), and also between group II and III (p=0.0012) (**Fig 4.B.vi**).

A significant increase (F_(1,12)_=112.7, p<0.0001) in the average diameter of altered and intact presynaptic mitochondria within the group I (p=0.0003), II (p<0.0001) and III rats (p=0.0002) was observed (**Fig 4.D.i**). The average area of mitochondria of the presynaptic terminal was significantly increased (F_(1,12)_=22.2, p=0.0003) between the altered and intact mitochondria in group I rats, and in group I (altered mitochondria) versus III (intact mitochondria) rats (**Fig 4.D.ii**). Significant change (F_(1,15)_=39.8, p<0.0001) in the circularity 18 index of altered and intact mitochondria within group I (p=0.004), II (p=0.03) and III (p=0.003) animals was also seen (**Fig 4.D.iii**).

### Synaptic vesicle

The synaptic vesicle density (SVs/nm^2^) in the NMJ of group II rats was significantly increased (F_(1,12)_=12.8, p=0.002) when compared to group I rats (p=0.0013), and also between group I and III (p=0.002) (**Fig 4.B.vii**). Docked synaptic vesicles, located within and beyond 200 nm from the plasma membrane were analyzed as vesicles located within the active zone (AZ); no significant difference was found in vesicle distribution within the AZ of Group I, II, and III rats (**Fig 4.B.viii**).

### Junctional folds

The number of junctional folds per synapse was significantly increased (χ^2^=11.9, p=0.0002) between group I and II rats (p=0.03) and between group II and III rats (p=0.004) (**Fig 4.C.i**). A significant increase in the average length of synaptic folds (χ^2^=19.5, p<0.0001) between group I and II (p<0.0001), and between group II and III rats (p=0.01) is observed (**Fig 4.C.ii**). The average junctional fold area (χ^2^=21.3, p<0.0001) was significantly increased in the group II to control (p<0.0001), and it was significantly decreased (p=0.02) in group III compared to II rats (**Fig 4.C.iii**). The opening width of junctional folds (χ^2^=15.4, p=0.0005) was significantly increased between group II compared to group I rats (p=0.0003), and also between control and recovery sleep group rats (p=0.05) (**Fig 4.C.iv**). The width of the synaptic cleft also showed significant alteration (χ^2^=15.4, p=0.0004), when compared between control and group II (p=0.002), and between group II and III animals (p=0.002; **Fig 4.C.v**).

### Acetylcholine and Acetylcholine esterase concentrations

The ACh concentration in the soleus muscle homogenate of the group II rats was significantly lowered compared to the control (χ^2^=7.7, p=0.007; **Fig 4.E.i**). When compared 19 between group I, II, and III rats, there was a significant increase (χ^2^=9.3, p=0.0005) in AChE in the soleus muscle homogenate in the group II (p=0.03) and III rats (p=0.03) compared to control (**Fig 4.F.i**).

### Correlation between neurotransmission and structural changes in NMJ of sleep deprived rats

In the group II rats, Spearman’s correlation between ACh and different morphometric variables revealed a strong negative correlation (r) with variables like average diameter of post-synaptic mitochondria (r=-0.8, p=0.33); average area of post-synaptic mitochondria (r=−0.6, p=0.41); density of pre-synaptic mitochondria (r=-1.0, p=0.08); width of synaptic cleft (r=-0.8, p=0.33). On contrary, ACh had positive correlation with density of post-synaptic mitochondria (r=0.8, p=0.33); opening width (r=0.8, p=0.33) of the NMJ of group II rats (**Fig 4.E.ii**). While AChE showed a strong negative correlation with density of post-synaptic mitochondria (r=-0.6; p=0.41), junctional fold opening width (r=-1.00; p=0.08) and positive correlation with variables like average diameter of post-synaptic mitochondria (r=0.60; p=0.41), average area of post-synaptic mitochondria (r=0.80; p=0.33), density of pre-synaptic mitochondria (r=0.80; p=0.33), width of synaptic cleft (r=0.60; p=0.41) of the 24 h sleep deprived rats (**Fig. 4.F.ii**).

## Discussion

Macrostructural examination of haematoxylin and eosin stained soleus muscle sections in rats subjected to 24 h sleep deprivation, reflected abnormality in fascicular architecture of myofibers compared to the two other groups. These changes were accompanied by mild variation in fiber size and myophagocytosis after 24 h sleep deprivation. Even after recovery sleep few atrophic and angulated fibers persisted. Previously, the endocrine correlates of muscle protein synthesis during sleep deprivation have been documented ^35^. The sleep deprivation protocol used in this study was similar to us, 20 where loss of muscle mass, cross sectional area of tibialis anterior muscle was reported, and it was associated with increase in corticosterone and decrease in testosterone of serum samples after sleep deprivation ^35^. Further, the authors have hypothesized that sleep restriction inhibits anabolic hormone secretion and facilitates catabolic hormones promoting protein degradation over synthesis causing muscle wasting ^36^. However, in these studies authors did not comment about the muscle hormone levels. Besides this, in our study, there was an increase in the area/field, perimeter and the density of the oxidative fibers (type I) of the soleus muscle in 24 h sleep deprived rats compared to the control. When the rats were allowed to recover from sleep deprivation, the area/field, perimeter and density of the type I fibers were restored as compared to the sleep deprivation. Souza et al., have shown that there were no differences in the area of myosin ATPase activity of gastrocnemius muscle type I fibers when compared between normal and 96 h REM sleep deprived rats, while a reduced area was observed in the type IIa fibers^37^. Moreover, 96 h sleep deprivation in male Wistar EPM-1 rats showed histopathological changes which confer that chonic sleep deprivation causes DNA damage, lipid peroxidation, and lysosomal activity prominently expressed in soleus muscle and not in plantar muscle. However, muscle mass was increased in soleus and reduced in plantar muscle of sleep deprived rats when compared to control ^38^. This observation suggests a differential effect depending upon the muscle type studied. This study was conducted in Wistar EPM-1 rats, and there were no sleep recovery group. The difference observed in our present study with the previous report could be due to the difference in the muscle type and the duration of sleep deprivation.

With the multiple platform method for sleep deprivation, we were able to increase the active and quiet wakefulness of the rats^27, 28^. When the rats were allowed to recover from sleep deprivation, AW time was significantly decreased, while the percentage of time spent in SWS and REM was significantly increased. Few brief sleep bouts were still observed during 21 sleep deprivation, which could be due to increased sleep drive in the rats making them fall asleep^39, 40^.

From our immunofluorescence data, it was observed that 24 h sleep deprivation caused amplification of presynaptic nerve terminal branching at the soleus muscle NMJ when compared to control rats. Similarly, the postsynaptic end-plate morphology along with the AChR area was increased in the NMJ of 24 h sleep deprived rats. These morphological changes were further supported by TEM findings, where the synaptic vesicle density, junctional fold morphology along with changes in a synaptic mitochondrial pool indicate that acute sleep deprivation for 24 h causes morphological changes at the peripheral synapse. Interestingly, in the 24 h sleep deprived rat, the biochemical estimation of ACh was lowered, while the concentration of its hydrolyzing enzyme (AChE) was higher. All these morphological and biochemical changes were reversed in rats allowed to recover from acute sleep deprivation. In a previous study by Gilestro et al., (2009), short period of sleep deprivation/ waking (6,12, 24 h) caused brain synaptic strengthening in the male *Drosophila***^5^**. In this study the association between sleep and synaptic plasticity was explored using multiple cue-based (viz. tactile, audio, and visual stimuli) sleep deprivation protocol. This deprivation protocol itself can be a cause for plastic changes found in the study. More so, though confocal microscopy was performed to confirm the specific brain region involved in the same, mainly the protein expression by Western blot of whole brain homogenates was used in this study^5^.

Differences in the size, appearance, and complexity of both pre and postsynaptic structures are observed from immunofluorescence. At the presynaptic level of 24 h sleep deprived rats, we have observed increased axonal diameters, axonal perimeters, increased branch number, total and average length per branch resulting in greater branching complexity. These were coupled with postsynaptic changes in the end-plate area, diameter, and perimeter lengths. The contribution of the skeletal muscle to NMJ formation during development and re-innervation has been reported previously^41^. Accordingly, the plastic changes of NMJs in the trained neuromuscular context include adaptations in endplate size, the sprouting of nerve terminal branches, and electrophysiological kinetics^42, 43^. These previous reports along with our findings point to activity-dependent changes in NMJ morphology and function.

Ultrastructural examinations of NMJ in our study revealed a significant increase in the 13 morphometric parameters from pre and postsynaptic junctional sites of 24 h sleep deprived rats. These include structures viz. mitochondria, synaptic vesicle, basal lamina, and junctional folds. These findings indicate that there is possible structural remodeling of NMJ in the acutely sleep deprived rats causing anatomical changes to combat 24 h sleep deprivation induced possible energy debt. This could be due to either diurnal rhythmicity of NMJ^44^, or due to activity-dependent plasticity^22, 42, 43^. In the study by Mehnert et al., the effect of light and dark phase on NMJ though tracing of motor neuron injected with HP in mutant *Drosophila* mutation in tim and per gene was observed. Morphological analysis showed that motor terminal morphology had a rhythmic pattern for day and night, which was linked with the clock genes ^44^.

We have also observed alteration in the pre and postsynaptic mitochondrial status at the NMJ, which is further correlated with the regulation of synaptic vesicle release^45, 46^. Altered presynaptic mitochondrial morphology was observed in the control and sleep deprived rats. Ruggiero et al., showed that the SW cycle affects the activity of certain mitochondrial enzymes which in turn regulate the mean firing rate of cortical and hippocampal neurons^47^. Inference from our TEM data further shed light on the link between the NMJ mitochondrial status and the SW cycle. Along with the synaptic mitochondrial morphology, there was also an increase in the synaptic vesicle density in the 24 h sleep deprived rats compared to the control.

Recently Cirelli and Tononi showed the effect of the SW cycle on the synaptic ultrastructure of the cerebrum and hippocampus. It was found though scanning electron microscopy that SW significantly influenced synaptic morphology which is correlated with the efficacy of AMPA receptors^3^. The expression of GluR1-containing AMPA receptors as an indicator of synaptic strength in the cortex and hippocampus was 30-40% higher after wakefulness in rats^10^. Similar to glutamate in the CNS, ACh is the principal neurotransmitter at the NMJ^45^. We have observed neuro-morphometric quantification of postsynaptic endplates of soleus muscles depicting significant changes in ACh area and perimeter, endplate area and perimeter, and in the number of ACh clusters after 24 h sleep deprivation when compared to control. Further, biochemical assessment of pre and postsynaptic neurotransmitters showed a reduction in the ACh and increased AChE concentration in the soleus muscle homogenate of 24 h sleep deprived rats. This shows the termination of synaptic transmission in the postsynapse conferring a functional upscaling of NMJ^45^. Moreover, we found a muscular adaptation to endurance after acute sleep deprivation though slow-twitch muscle conversion. Ultra-structural alterations in the increased area of the junctional fold and the width of the synaptic cleft are also evident from our TEM data. Increased junctional fold area increases muscle surface area allowing to hold more ACh^45^. Moreover, we have observed widening of the synaptic cleft, which is filled with extracellular matrix called synaptic basal lamina containing AChE^45, 48^. These correlated structural findings with biochemical quantification of AChE, imply increased demand of neuromuscular transmission at the NMJ during 24 h sleep deprivation.

With the reduced consumption of energy by synaptic transmission during hyperpolarized down states, SWS represents an elective time for brain cells to carry out many housekeeping functions, including protein translation, the replenishment of calcium in presynaptic pools, the replenishment of glutamate vesicles, the recycling of membranes, the resting of mitochondria^2, 49–51^, and the metabolite clearance from the extracellular space^52^. Recently we have reported that the muscle temperature was least altered during the normal SW cycle in rats, indicating that probably the muscle atonia during REM sleep provides a conducive environment for the muscle to rest and repair^14^, somewhat similar to slow-wave activity during SWS for CNS synaptic remodeling.

The correlation between ACh with synaptic morphological variables in our study suggests that acute sleep deprivation for 24 h causes neuromuscular changes in synaptic energy homeostasis though pre and postsynaptic mitochondrial modification respectively for vesicular transport and postsynaptic currents^46, 53^. There was an additional change in the alignment of two synaptic compartments via thickening of the basal lamina to compensate for the loss of ACh in soleus muscle in the 24 h sleep deprived rats. The correlation between AChE with various NMJ structures reveal that the pre and postsynaptic homeostasis is altered along with junctional matrix alignment leading to the expression of more AChE, which terminates synaptic transmission by hydrolyzing ACh^45^. Furthermore, correlating the morphometric variables with neurotransmitter concentration also exhibited a similar heat-map revealing a structural remodeling in the NMJ after acute sleep deprivation, which is probably due to the energy balance in the synapse closely associated with the neuromuscular transmission^54^.

The NMJ is a complex structure that mediates the cross-talk between motor neurons and muscle fibers^23, 45^. Muscle is the other excitable tissue besides neurons and the NMJ is the most studied structure for synaptic physiology. Effective neurotransmission depends upon the regular arrangement of postsynaptic AChs and proper alignment between the terminal bouton and underlying motor endplate^23^. The outcome of our study suggests substantial remodeling 25 of the NMJ during acute sleep deprivation and after recovery sleep. These changes reflect plasticity at both pre and postsynaptic structures of NMJ, and indicate at possible correlation to neuromuscular transmission and muscle function. Our findings provide strong evidence for the first time on the influence of sleep on the muscle and neuromuscular junction morphology and functions.

## Data Availability

Details of materials and experimental protocols, the sources and catalogue numbers of reagents necessary for replication of the study are included in the manuscript. All raw data are deposited in the centre for open science and can be accessed from the following link: (https://mfr.osf.io/render?url=https://osf.io/f29n3/?direct%26mode=render%26action=download%26mode=render).

## Acknowledgements

This work was supported by the All India Institute of Medical Sciences, New Delhi and the Indian Council of Medical Research, New Delhi (45/6/2019/PHY/BMS). We want to extend our acknowledgement to Dr. M.C Sharma and Dr Soumya Sahu from Department of Pathology, and Dr. Shivam Pandey, Department of Biostatistics, All India Institute of Medical Sciences, New Delhi for re-evaluation of histopathological data and statistical results respectively.

## Author contributions

B.S., A.R., H.N.M., N.A., and R.N. designed research; B.S., A.R., L.C. performed research; B.S., A.R and T.S.G. analyzed data; A.S., T.C.N., contributed to transmission electron microscopy facility; B.S., A.R and T.S.G. wrote the paper; and B.S., A.R., T.S.G., H.N.M., performed critical drafting of the manuscript.

